# LIME: a fully automated pipeline for high-throughput quantification of leaf lesions

**DOI:** 10.64898/2026.05.07.723432

**Authors:** Xavier Schneider, Da Tan, Lara Carroll, Rhey Caners, Mark Belmonte, Steve Whyard, Christopher J. Henry

## Abstract

Accurate quantification of leaf lesion severity is essential for plant disease research and phenotyping but is often limited by subjective visual scoring and time-intensive manual image analysis. We present LIME, a fully automated, open-source image analysis pipeline for high-throughput quantification of leaf lesions from disease assay images. LIME integrates zero-shot leaf segmentation using the Segment Anything Model with a convolutional neural network for lesion area estimation.

Applied to *Arabidopsis thaliana* leaves infected with *Sclerotinia sclerotiorum*, the proposed approach achieved a mean absolute percentage error of 12.9%, comparable to observed intrarater variability in manual scoring. Stratified evaluation across lesion-size groups demonstrated consistent prediction accuracy for small, intermediate, and large lesions, and comparative analysis showed that the deep learning–based model substantially outperformed color-based baseline methods. Under GPU-accelerated execution, LIME processed complete assays containing approximately 200 leaves in 15 minutes, representing an approximate 13-fold reduction in processing time relative to manual annotation.

Together, these results indicate that LIME enables objective, reproducible, and scalable quantification of leaf lesion severity in standardized plant pathology assays. The pipeline is released as an open-source tool to support quantitative phenotyping studies.

## Introduction

Plants are frequently attacked by pathogenic microorganisms in both natural and agricultural systems, leading to observable disease symptoms such as chlorotic or necrotic leaf lesions. Accurate quantification of disease severity is essential for several reasons: i) it enables the linkage between disease progression and yield loss; ii) it supports the evaluation of germplasm and cultivars in plant breeding programs; iii) it informs disease management strategies such as targeted pesticide application during epidemics [1]. In addition, quantitative disease assessment contributes to a deeper understanding of fundamental biological processes, including host–pathogen coevolution and plant disease epidemiology [1].

Disease severity and host resistance can be assessed using a variety of approaches, including measurements of bacterial load [2] and fungal load [3], detection of reactive oxygen species (ROS) [4], quantification of marker gene expression [5], and visual assessment of leaf symptoms [6]. Among these methods, visual assessment of leaf symptoms remains particularly attractive due to its conceptual simplicity, minimal training requirements, and relatively high throughput [6]. Depending on the pathogen and host, symptoms may manifest as chlorosis, as observed in *Arabidopsis thaliana* infected with *Pseudomonas syringae* pv. *tomato* DC3000, or as necrotic lesions, such as those caused by necrotrophic fungi including *Sclerotinia sclerotiorum* and *Botrytis cinerea*.

Historically, macroscopic disease symptoms have often been evaluated using ordinal rating scales, or disease indices, which categorize symptom severity from minimal to severe [1]. While widely adopted, these assessments rely heavily on human judgment and are therefore subject to observer bias, interrater and intrarater variability, and limited scalability. As a result, there is growing demand for objective, quantitative, and reproducible methods for assessing disease symptoms at scale [7].

On the other hand, once individual leaves have been identified and isolated from assay images, a subsequent task is to quantify disease severity. In plant pathology, severity is most commonly assessed by measuring the proportion of leaf area exhibiting visible disease symptoms [8]. Even when performed by experienced specialists, this process is challenging, as evaluators must visually interpret lesion boundaries and align observations with diagnostic guidelines. As a result, manual assessments inherently involve subjectivity, and the resulting annotations cannot be regarded as absolute ground truth [8]. This subjectivity gives rise to both intrarater variability, reflecting differences when the same annotator repeats measurements, and interrater variability, reflecting disagreement across annotators.

To meet this demand, image-based approaches have become increasingly common for quantifying fungal lesions, particularly those involving necrotic tissue. Manual or semi-automated workflows based on tools such as ImageJ remain widely used [9, 10]. Although these tools enable pixel-based lesion area estimation and diameter measurements, they are time-intensive and often struggle to robustly quantify irregular lesion morphologies [6]. Alternative solutions, including dedicated hardware such as the YMJ-C smart leaf area meter [11] and commercial software such as Adobe Photoshop [12], can partially alleviate these challenges but frequently incur higher costs or require substantial manual intervention [6].

More recently, several image analysis tools have been developed to streamline disease quantification. Examples include PIDIQ [13], which focuses on pathogen-induced chlorosis, Colour-analyzer [6], a web-based platform for fungal lesion quantification, and Sc-Analyzer [7], which employs a flatbed scanner and grid-based layout to estimate pathogen spread. However, many existing tools remain semi-automated, require specialized imaging equipment, or depend on manually tuned color thresholds, limiting their robustness and reproducibility across experimental conditions [6].

In this context, there is a clear need for fully automated, reproducible pipelines that can robustly quantify leaf lesions across large-scale assays while minimizing human intervention. In this study, we focus on lesions caused by the necrotrophic fungus *Sclerotinia sclerotiorum* in *Arabidopsis thaliana*, a pathosystem commonly used to study host resistance mechanisms. This work supports broader efforts to develop transgenic plants with enhanced resistance to *S. sclerotiorum*, including strategies based on RNA interference targeting essential fungal genes or virulence factors [14].

Here, we introduce the *Lesion Inference & Measurement Engine* (LIME), an open-source, fully automated command-line tool for high-throughput quantification of lesion formation on plant leaves. LIME integrates a foundation-model-based segmentation stage with a deep learning regression model to estimate lesion area and total leaf area, enabling precise calculation of infected tissue proportions across hundreds of samples per assay. By eliminating manual thresholding and subjective scoring, LIME substantially reduces observer-dependent variability while providing throughput suitable for large-scale phenotyping studies.

The main contributions of this work are as follows: i) We present a fully automated, end-to-end pipeline for quantitative leaf lesion measurement that requires no manual annotation or threshold tuning during inference. ii) We demonstrate the use of a zero-shot segmentation foundation model to robustly isolate individual leaves from high-density assay images under variable lighting conditions. iii) We introduce a deep learning–based lesion area estimation model that achieves accuracy comparable to human intrarater variability while providing substantial gains in throughput. iv) We release LIME as an open-source, reproducible tool designed for integration into standardized plant pathology workflows.

## Related work

### Traditional lesion quantification methods

Traditional approaches for assessing foliar disease severity, including visual rating scales and manual or semi-automated image analysis tools, are widely used in plant pathology and phenotyping. These methods, along with their limitations in terms of subjectivity, scalability, and dependence on manual parameter tuning, are discussed in detail in the Introduction. While these approaches reduce reliance on purely visual scoring, they typically require manual interaction, fixed thresholds, or specialized acquisition setups, which can limit throughput and reproducibility.

### Segmentation methods in plant phenotyping

Traditional segmentation approaches in vision-based plant phenotyping have largely relied on classical image processing techniques. ApLeaf [15], for example, employs grayscale thresholding to group pixels by intensity and separate foliage from background. Otsu’s method [16] automatically determines an optimal threshold by minimizing within-class variance and has been widely adopted for background subtraction in automated plant recognition systems [17, 18]. Another commonly used technique is the watershed transformation, which interprets an image as a topographic surface and segments regions by flooding from predefined seed points until strong gradient boundaries are encountered [19].

Despite their simplicity and efficiency, these methods are highly sensitive to imaging conditions. Variations in illumination intensity, angle, or spectral composition can substantially alter leaf appearance [20, 21], leading to inconsistent contrast, poorly defined boundaries, and reduced segmentation accuracy. Shadows, glare, and uneven lighting further complicate reproducible leaf delineation, limiting the robustness of both automated and semi-manual segmentation pipelines [20, 21].

Deep learning–based segmentation methods have demonstrated improved robustness under such variability. Convolutional neural network architectures such as U-Net [22] and DeepLab [23] learn hierarchical feature representations from annotated data and can reliably separate plant structures from background across a wide range of imaging conditions. However, the effectiveness of supervised segmentation models depends heavily on the availability of large, pixel-level annotated datasets, which are costly to produce and often unavailable for specialized plant pathology assays [24].

To mitigate this limitation, recent work has explored the use of large foundation models for segmentation. In this study, we leverage the Segment Anything Model (SAM) [25], a promptable segmentation model pretrained on the SA-1B dataset comprising over 11 million images and one billion masks. SAM provides strong zero-shot segmentation performance across diverse visual domains, making it well suited for scenarios where domain-specific annotated training data are limited. Rather than relying on fully supervised retraining, SAM is used here to generate candidate leaf masks that are subsequently refined using domain-specific constraints.

### Computational approaches for leaf lesion assessment

A variety of computational approaches have been proposed for automated disease spot detection and lesion quantification. Early image-processing methods often relied on color space transformations and handcrafted rules. Patil and Bodhe convert RGB images to the HSI color space and apply triangle-based thresholding to isolate diseased regions [26]. Pagola et al. [27] quantify nitrogen deficiency in barley leaves by manipulating RGB channels and applying principal component analysis (PCA) to derive pixel-wise greenness indices. Other approaches include fuzzy logic–based systems [28] and rule-based, knowledge-driven frameworks [29].

More recently, machine learning and deep learning methods have become increasingly prevalent for plant disease detection and quantification [24]. These approaches offer improved flexibility and robustness relative to handcrafted pipelines, particularly in the presence of variable lesion morphology, illumination conditions, and background complexity. Consequently, LIME adopts a deep learning–based framework for lesion severity estimation, enabling direct prediction of lesion area from image data while reducing reliance on manually defined thresholds or heuristics.

## Materials and methods

### LIME pipeline overview

LIME is an automated image analysis pipeline for quantifying leaf lesions from high-resolution assay images. Given a single image containing multiple leaves arranged on a uniform background, the pipeline produces per-leaf measurements of total leaf area and lesion area. An overview of the workflow is shown in Fig. 1.

**Fig 1.**
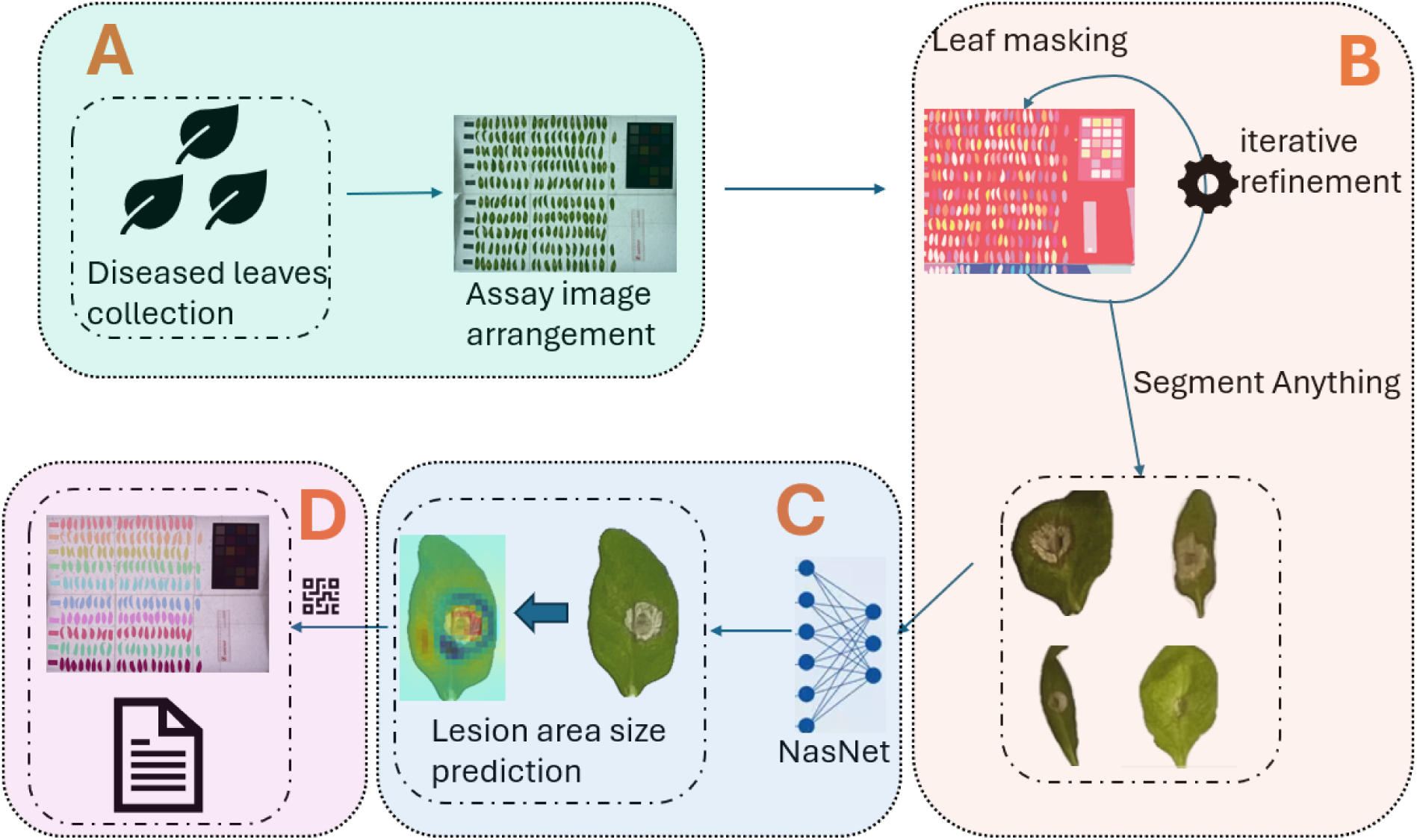
Overview of the LIME workflow. Raw assay images are processed through four stages: (A) fungal infection assay and collection of diseased leaves, (B) zero-shot generation of candidate leaf masks and mask refinement to isolate individual leaves, (C) per-leaf lesion area estimation using a convolutional neural network, and (D) group-wise results collection using QR code recognition.

The pipeline consists of four sequential stages. First, detached leaves exhibiting disease symptoms are collected and arranged on a uniform background to form a standardized assay image (Fig. 1A). Second, candidate segmentation masks are generated directly from the raw assay image using a pretrained, zero-shot segmentation model to identify potential leaf regions without manual prompts or annotations. These candidate masks are refined through an automated filtering process to remove background elements and produce a set of non-overlapping masks corresponding to individual leaves (Fig. 1B). Third, each segmented leaf is analyzed independently by a convolutional neural network to estimate lesion area, yielding per-leaf quantitative measurements suitable for downstream analysis (Fig. 1C). Finally, metadata, such as QR-code-based sample identifiers, can be incorporated during post-processing to associate measurements with experimental conditions (Fig. 1D). The following sections describe each stage of the pipeline in detail.

### Assay setup and image acquisition

Leaf lesion assays involved detached Sclerotinia-infected *Arabidopsis thaliana* leaves arranged on a uniform white background. The leaf infection experiment follows the methods of [3]. Each assay consisted of approximately 200 individual leaves placed in rows with minimal overlap between neighboring samples. Leaves were positioned to ensure clear visual separation and consistent orientation across the assay image; the color palette is a predefined set of RGB colors used to assign each mask group a distinct, consistent overlay color on the output image so humans can easily tell groups apart; the ruler is used for physical scaling when calculating the sizes of the lesion area. (Fig. 2A).

**Fig 2.**
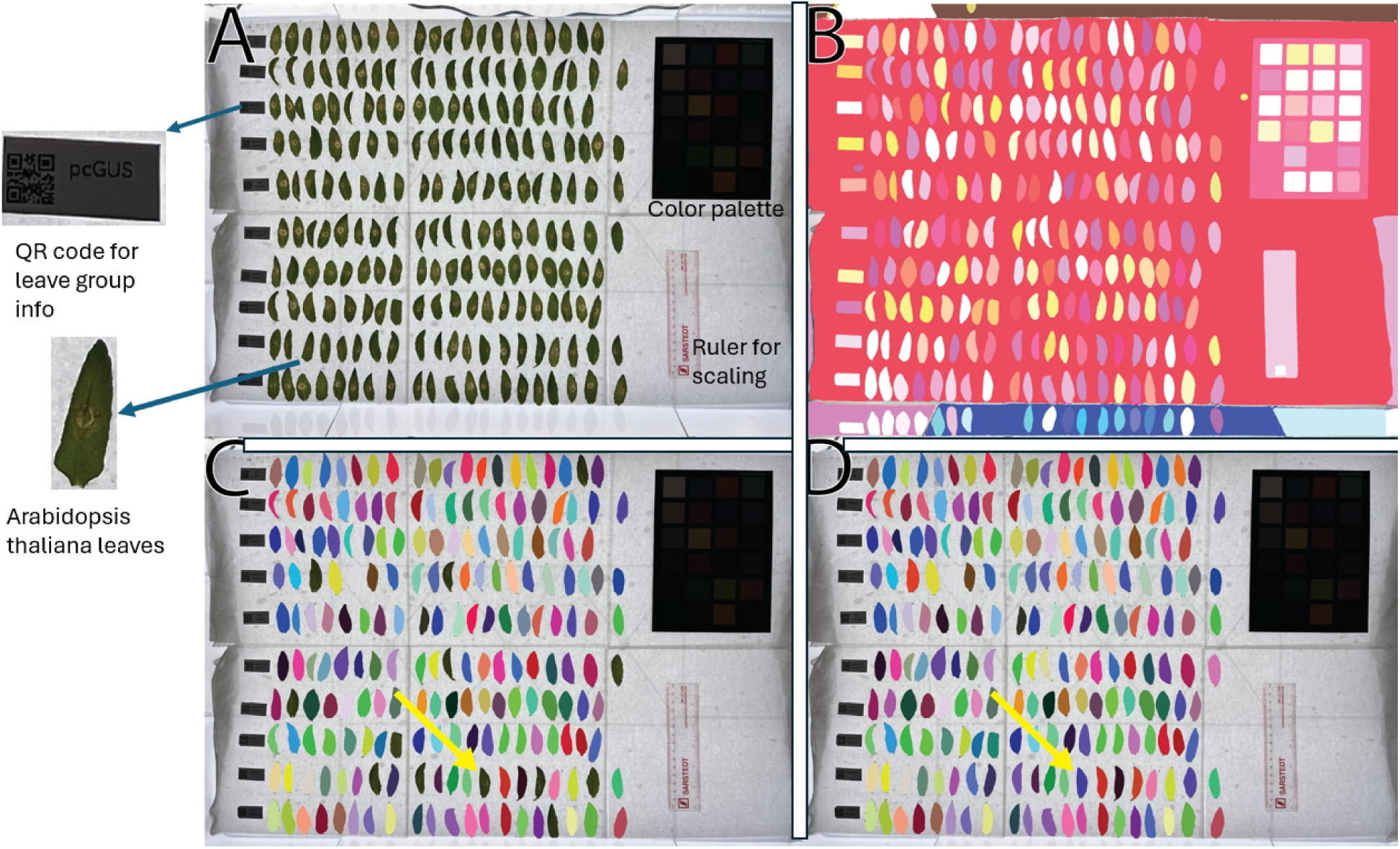
Iterative refinement of assay-level leaf segmentation. (A) Assay image containing approximately 200 *Arabidopsis thaliana* leaves infected with *Sclerotinia sclerotiorum*. (B) Assay image overlaid with candidate segmentation masks generated by SAM. (C) Assay image overlaid with masks after one iteration of the refinement algorithm. (D) Assay image overlaid with masks after two iterations of refinement. The mask indicated by the yellow arrow is correctly classified as a leaf after refinement.

Images were acquired using a digital camera mounted at a fixed height above the assay surface to maintain consistent imaging geometry across experiments. Camera position, focal length, and lighting conditions were kept constant within and across assays. A physical scale (ruler) was included within the field of view of each image to enable conversion from pixel-based measurements to physical units (Fig. 2 A).

For experiments involving multiple genotypes or experimental conditions, leaves corresponding to each group were arranged in separate horizontal rows. A QR code encoding the sample identifier was placed at the beginning of each row. These QR codes were later used during post-processing to associate segmented leaves with their corresponding experimental groups.

All images were captured at high resolution and stored in a lossless format prior to analysis. No manual cropping, thresholding, or image enhancement was performed before processing with LIME. The raw assay images served as the sole input to the pipeline.

### Leaf segmentation

#### SAM-based mask generation

The first step of leaf segmentation in LIME is the generation of candidate object masks from the raw assay image. For this purpose, we employ the Segment Anything Model (SAM) [25], a promptable segmentation model pretrained on a large-scale, diverse image corpus.

Given an input assay image, SAM is applied in a fully automatic manner to produce a set of candidate segmentation masks (Fig. 2 B). Each mask corresponds to a contiguous region of pixels that SAM identifies as a potential object. No user-defined prompts, bounding boxes, or point annotations are provided during inference. As a result, the output of this stage consists of a collection of overlapping and non-overlapping masks representing both leaf and non-leaf regions within the image.

The generated masks are retained in their original resolution and treated as candidate regions for subsequent processing. At this stage, no assumptions are made regarding object class, and no filtering or thresholding is applied. The purpose of this stage is solely to enumerate potential leaf regions with high recall, deferring all selection and refinement decisions to the subsequent mask refinement step.

#### Iterative mask refinement

The candidate masks produced by SAM include both leaf and non-leaf regions and therefore require further refinement to isolate individual leaves. LIME performs this refinement through an iterative, color-based filtering procedure that progressively identifies masks corresponding to leaf tissue while rejecting background elements and spurious regions (Alg. 1).

At the start of the refinement process, a representative leaf mask is selected from the set of candidate masks to initialize a reference color profile. The color distribution of a mask *m* is computed in the hue–saturation–value (HSV) color space and represented as a normalized histogram

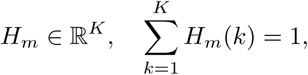

where *K* denotes the number of histogram bins. The initial reference histogram is denoted by *H_r_* (Fig. 3).

**Fig 3.**
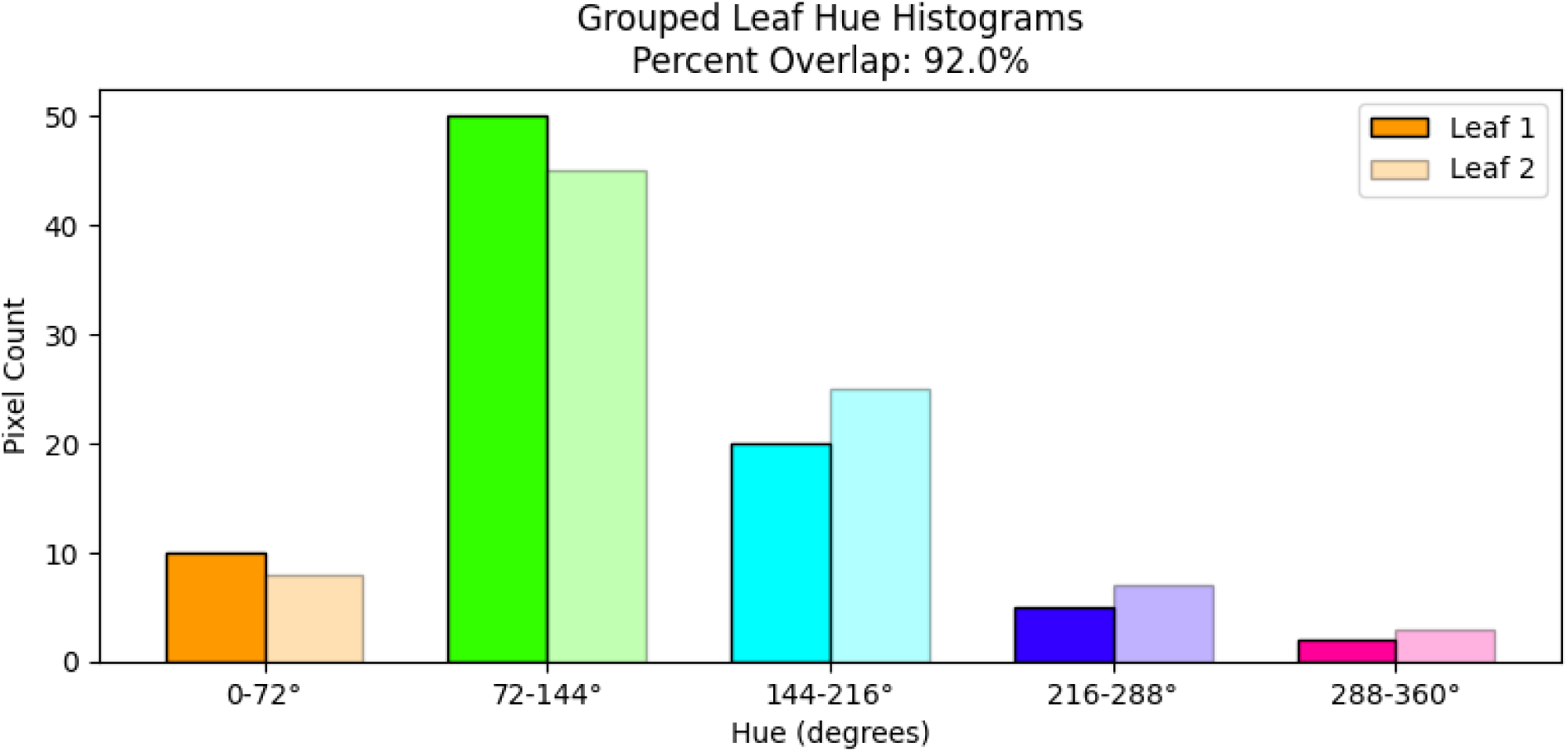
Reference-profile comparison during mask refinement. Comparison of a candidate mask (Leaf 2) and the reference profile (Leaf 1) used in the iterative refinement algorithm.

For each remaining candidate mask *m*, an HSV histogram *H_m_* is computed and compared to the current reference profile using histogram intersection,

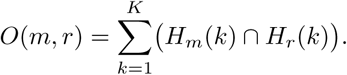

A candidate mask is provisionally classified as a leaf if *O*(*m, r*) *> τ*, where *τ* is a fixed threshold for overlapping (10%).

To account for natural variation in leaf coloration and illumination conditions, the reference histogram is updated iteratively. After each iteration *t*, the reference profile is updated by averaging the histograms of newly accepted leaf masks *A^(t)^*,

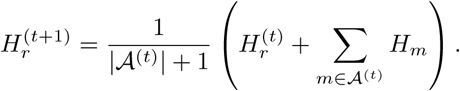

In parallel, geometric and topological constraints are applied to exclude implausible leaf candidates. Masks are discarded if they contain multiple disconnected components, exhibit extreme concavity as measured by low solidity, or overlap substantially with previously accepted leaf masks. The refinement process is repeated until the color profile of the reference leaf stabilizes, defined as an iteration in which no additional candidate masks satisfy the acceptance criteria (Fig. 2 C,D).

The final output of this stage is a set of non-overlapping binary masks, each corresponding to an individual leaf and suitable for downstream lesion analysis.

### Lesion area estimation

Following leaf segmentation, LIME estimates lesion area for each individual leaf using a convolutional neural network. This stage operates on cropped leaf images derived from the segmentation masks and produces a single scalar value corresponding to the predicted lesion area.

#### CNN architecture

To estimate lesion size on each leaf, we adopt a modified NASNet A backbone. NASNet A—introduced by Zoph et al. [30] via Neural Architecture Search (NAS)—uses reinforcement learning to assemble two types of modular “cells”: normal cells, which preserve spatial dimensions for feature extraction, and reduction cells, which downsample feature maps while increasing their depth. Each cell is defined as a directed acyclic graph combining operations like depthwise separable convolutions, pooling layers, and skip connections. Thanks to its automated search process, NASNet A scales gracefully: cell structures optimized on small datasets (e.g., CIFAR 10) transfer effectively to large benchmarks such as ImageNet. In our pipeline, we load the ImageNet pretrained NASNet A, replace its classification head with a regression head, and train end-to-end using a mean absolute percentage error (MAPE) loss to predict per leaf lesion area.

#### Training and inference

The dataset used in this study consists of 1,700 images of detached *Arabidopsis thaliana* leaves infected with *Sclerotinia sclerotiorum*. Individual leaves were imaged following standardized lesion assays and subsequently segmented and cropped to a fixed spatial resolution of 225 × 225 pixels for lesion analysis. Ground-truth lesion area annotations were generated using ImageJ (note that rater variability was measured at 10% MAPE). For each leaf image, lesion regions were manually delineated and quantified in pixel units. These pixel-based lesion area measurements served as reference labels for training and evaluating the lesion area estimation model. The dataset was randomly divided into training, validation, and test sets using a 70% / 15% / 15% split. All splits were performed at the leaf level, and no images were shared across subsets.

Lesion area estimation was formulated as a regression problem: given a cropped leaf image *x_i_*, the network predicted a scalar lesion area *y*^*_i_* = *f_θ_*(*x_i_*), which was compared against the ground-truth lesion area *y_i_*. Model parameters *θ* were optimized by minimizing the mean absolute percentage error (MAPE), defined as

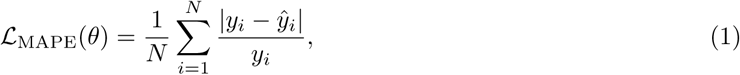

where *N* is the number of training samples. This objective penalizes the relative error between predicted and true lesion sizes, helping balance contributions from leaves with small and large lesions.

During inference, each segmented leaf was processed independently to produce a lesion area estimate,

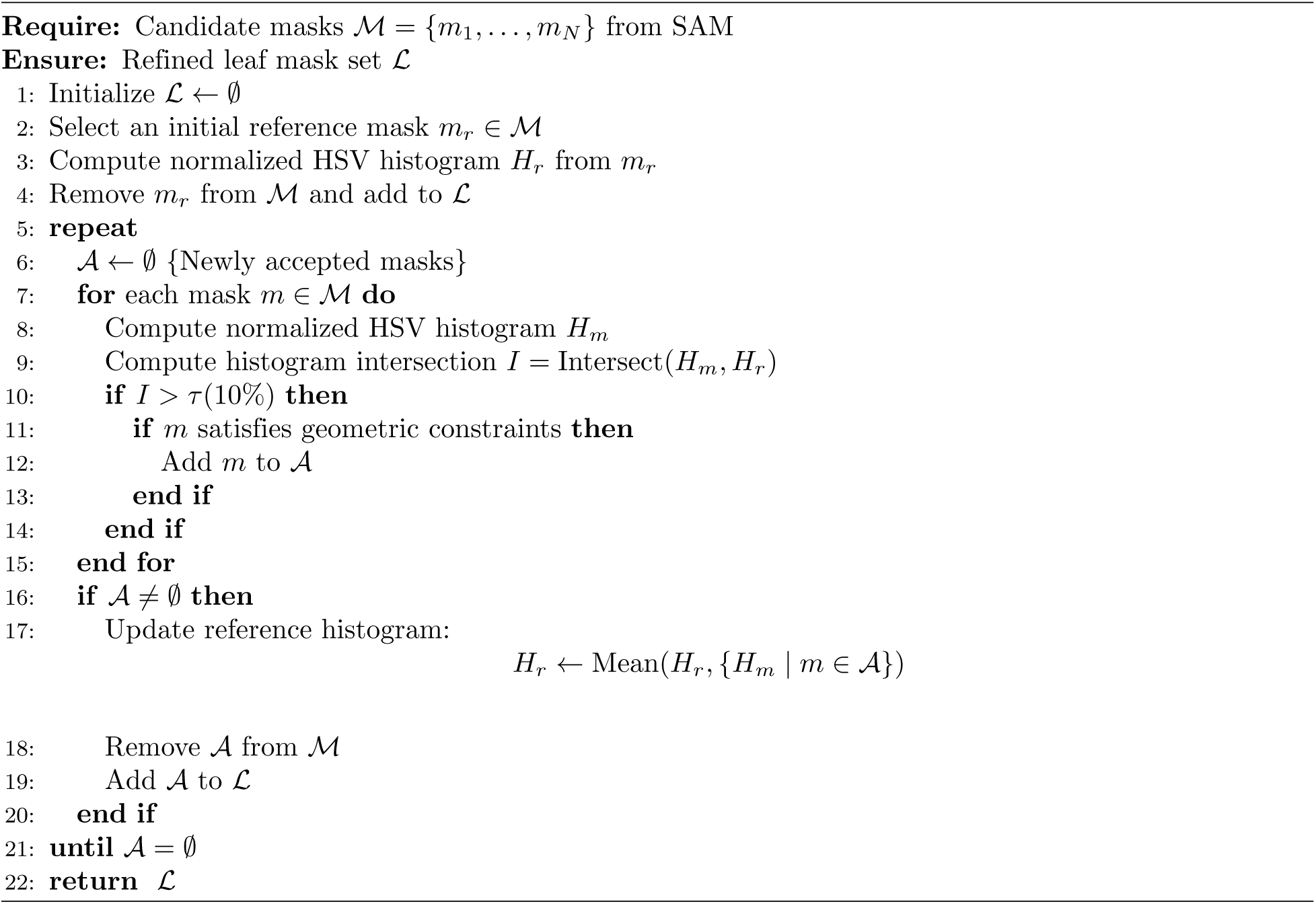
Algorithm 1 Iterative leaf mask refinement.

which was retained in pixel units and passed to the post-processing stage.

### Post-processing and data export

Following lesion area estimation, LIME performs a series of post-processing steps to convert pixel-based measurements into physical units, associate individual leaves with experimental metadata, and export results in structured formats.

Total leaf area is computed for each segmented leaf by summing the pixels within its binary segmentation mask. Lesion area predictions produced by the regression model are also expressed in pixel units. To convert both measurements to physical units (e.g., mm^2^ or cm^2^), a calibration factor is computed from the in-frame physical scale included in each assay image. This factor represents the physical area per pixel and is applied uniformly across all measurements within the image.

Our assays contain multiple experimental groups, and leaves are arranged in horizontal rows, each labeled with a QR code encoding the corresponding sample identifier. QR codes are detected by LIME and decoded from the assay image and used to identify the row associated with each experimental condition. Individual leaf masks are assigned to rows by clustering their centroid *y*-coordinates using a Gaussian Mixture Model, with the number of clusters determined by the number of detected QR codes. This procedure enables automatic association of each leaf with its corresponding experimental group.

Quantitative results are exported as comma-separated value (CSV) files, containing per-leaf measurements of total leaf area, lesion area, and derived disease severity metrics. In addition, LIME generates an annotated version of the original assay image in which each segmented leaf is outlined and labeled, enabling easy cross reference between the CSV entries and the physical leaves in the assay (Fig. 4).

**Fig 4.**
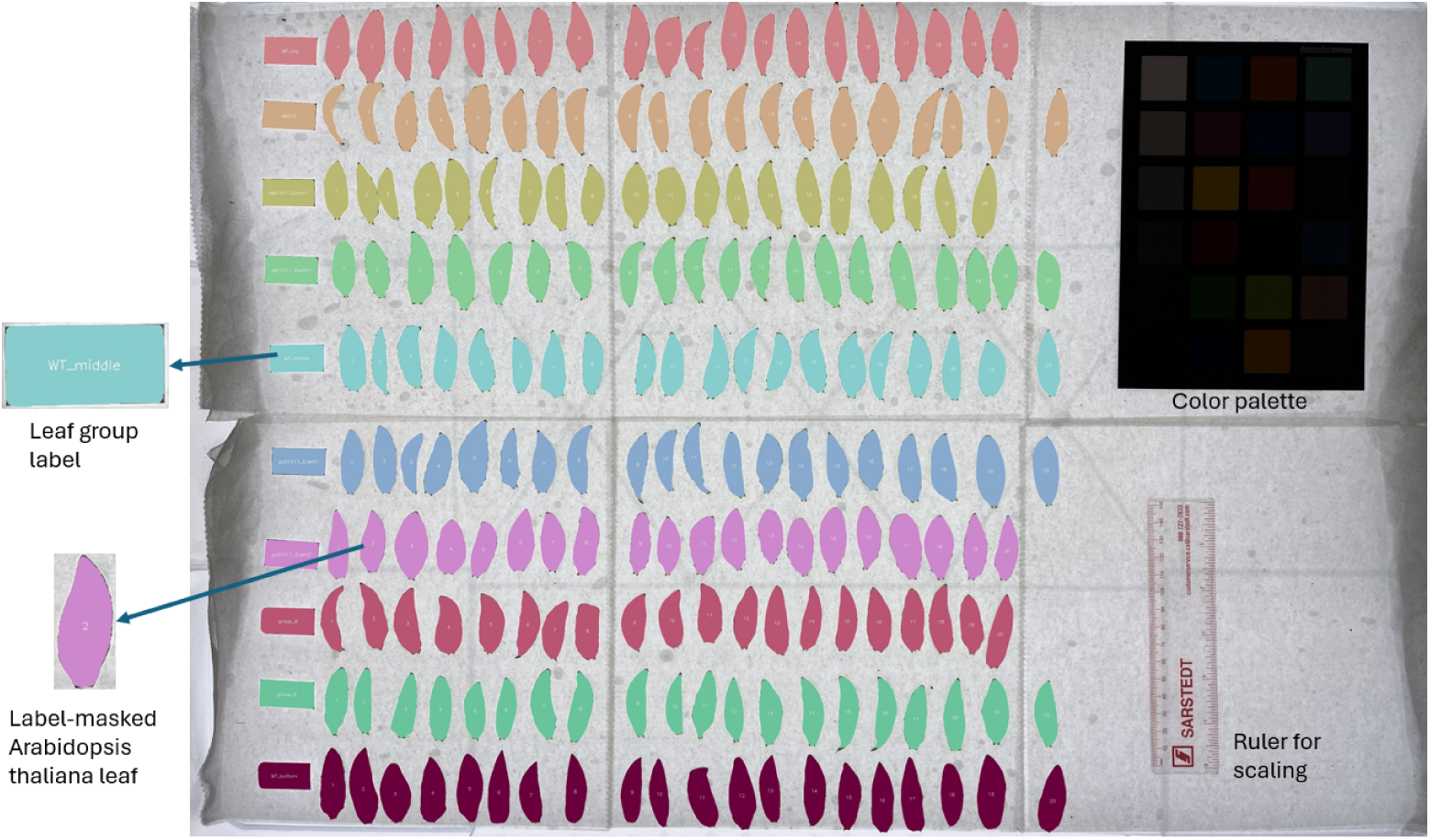
QR code-based labeling of segmented leaves. Segmented leaves are sorted, labeled, and assigned to experimental groups using QR code metadata.

These outputs are the final result of the pipeline and are used for downstream statistical analysis and visualization.

### Color-based baseline methods

To provide interpretable and lightweight baselines for lesion quantification, we implemented two color-based lesion detection methods operating in the HSV color space. Both baselines estimate lesion area by identifying healthy leaf tissue and treating remaining leaf pixels as lesions. These approaches do not require learning during inference and rely only on color statistics derived from manually sampled data.

#### HSV threshold-based lesion detection

The first baseline uses fixed HSV thresholds to separate healthy leaf tissue from lesion regions. Input leaf images are first converted from RGB to HSV color space. Pixels corresponding to the white background are removed using a high-value, low-saturation threshold. All remaining pixels are treated as leaf tissue.

Healthy leaf pixels are identified using empirically derived HSV thresholds obtained from manual color sampling of lesion and healthy regions. The samples are from 30 random leaf images from the training dataset. Fig. 5 shows the distribution of the hue, saturation and value levels of the sampled pixels. Based on these samples, healthy tissue is characterized by a narrow hue range corresponding to green leaf color, high saturation values, and moderate value intensity. Pixels falling within this predefined healthy range are classified as healthy leaf tissue. All other non-background pixels are classified as lesion candidates.

**Fig 5.**
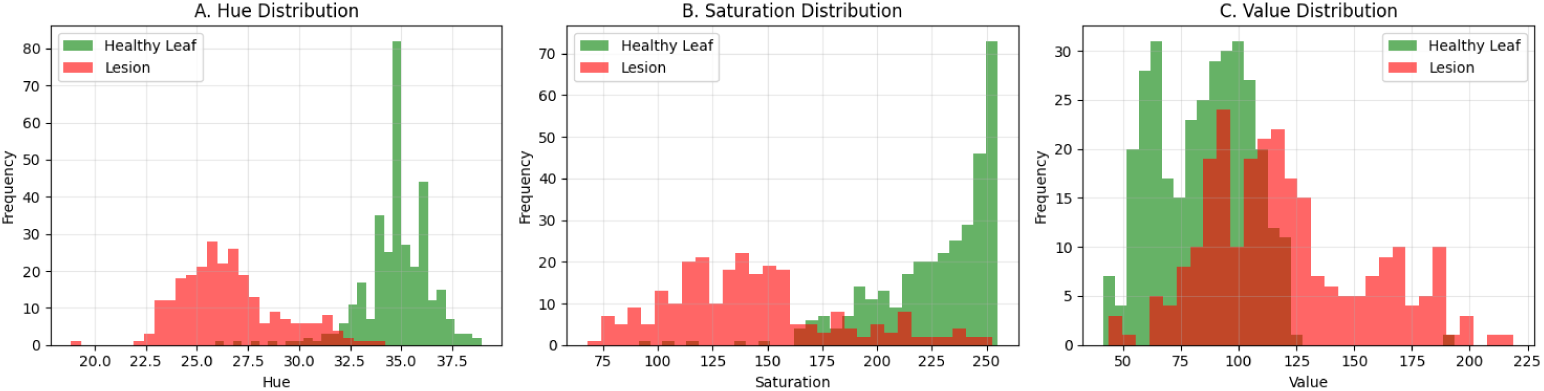
HSV distributions of sampled leaf pixels. Panels (A)–(C) show the distributions of hue, saturation, and value, respectively, for sampled *Arabidopsis thaliana* leaf pixels from the training dataset. Healthy and lesion regions exhibit distinct distributions in hue (A) and saturation (B).

To reduce noise and spurious detections, morphological opening and closing operations are applied to the lesion mask using a small structuring element. Connected components smaller than a minimum area threshold are subsequently removed. The final lesion area is computed as the number of pixels classified as lesion.

#### K-means clustering-based lesion detection

The second baseline employs unsupervised clustering to partition leaf pixels into color-consistent groups. After conversion to HSV color space, background pixels are excluded using the same white-background criterion as in the threshold-based method. The remaining pixels are clustered in HSV space using K-means clustering with a fixed number of clusters.

Cluster centers are analyzed to identify the cluster corresponding to healthy leaf tissue. This identification is based on learned color characteristics of healthy leaves, namely high saturation and hue values within the expected green range. A composite score combining saturation magnitude and proximity of hue to the healthy-leaf mean is used to select the healthy cluster.

All pixels not assigned to the healthy cluster are classified as lesion pixels. Lesion area is computed as the total number of pixels assigned to non-healthy clusters. No spatial smoothing or post-clustering morphological filtering is applied in this baseline.

#### Area conversion and batch inference

For both color-based baselines, lesion area is initially computed in pixel units for each individual leaf image. Pixel-based lesion areas are converted to physical units using the same image calibration factor applied in the main LIME pipeline. Baseline inference is performed independently for each leaf image, enabling direct comparison with the NASNet–based lesion estimation results. These baselines were evaluated using the same dataset, metrics, and calibration procedure as the LIME pipeline.

### Evaluation metrics

Model performance for lesion area estimation was evaluated using the MAPE, defined in Equation 1. MAPE was selected to provide a scale-normalized measure of prediction error across leaves exhibiting a wide range of lesion sizes. Reported performance values correspond to evaluation on the held-out test set. In addition, we also reported mean absolute error (MAE) for the model’s performance.

### Hardware and runtime configuration

All experiments were conducted using a workstation equipped with an NVIDIA RTX 3060 Ti GPU. GPU acceleration was used for both segmentation and lesion area estimation stages. While LIME can be executed on CPU-only systems, GPU acceleration is recommended due to the computational cost of segmentation models.

Under this configuration, LIME processed a complete assay image containing approximately 200 leaves in approximately 15 minutes, including segmentation, lesion estimation, and post-processing steps. For comparison, manual lesion scoring required approximately 60 seconds per leaf, resulting in a total processing time of roughly 3 hours for an equivalent assay.

### Baseline methods and evaluation protocol

In addition to the deep learning–based lesion estimation model, two color-based baseline methods were implemented to provide interpretable reference points for comparison. As mentioned in Section 4.5, the baselines consist of (i) an HSV threshold-based method and (ii) a K-means clustering–based method operating in HSV color space. Both approaches estimate lesion area by identifying healthy leaf tissue and classifying remaining leaf pixels as lesions, without the use of learned models during inference.

Baseline inference was performed on the same set of individual leaf images used for evaluating the lesion estimation model. Input images, background removal, and pixel-to-physical area conversion were identical across all methods to ensure a fair comparison. For each leaf, lesion area was computed in pixel units and converted to physical units using the same calibration factor derived from the assay image.

Performance of the baseline methods was evaluated using the same metrics applied to the deep learning–based approach, including MAPE, and MAE. Ground-truth lesion areas were obtained from manual ImageJ annotations.

### Lesion-size group stratification

In addition to overall performance evaluation, lesion area predictions were analyzed across lesion-size groups to assess model behavior under different disease severities. Lesion-size stratification was performed on the held-out test set using ground-truth lesion area annotations of 255 leaves.

Test samples were divided into three lesion-size groups, small, intermediate, and large, based on fixed lesion area thresholds derived from the empirical distribution of lesion sizes in the test dataset. Thresholds were selected such that each group contained an equal number of samples, enabling balanced comparison across severity levels. Lesion area values are reported in pixel units.

The resulting lesion-size group definitions and sample distribution for the test set are summarized in Table 1. All evaluated methods, including the deep learning–based model and the two color-based baselines, were assessed independently within each lesion-size group using the same evaluation metrics described in the Evaluation metrics subsection.

**Table 1.** Lesion-size group definitions. Sample distribution and lesion-area ranges for the held-out test set. Lesion area is reported in pixel units.

| Lesion-size group | Lesion area range (pixels) | # samples |
| --- | --- | --- |
| Small | < 1656.96 | 85 |
| Intermediate | 1656.96 – < 3024.38 | 85 |
| Large | ≥ 3024.38 | 85 |

## Results

### Segmentation robustness (qualitative)

The leaf segmentation stage of LIME was evaluated qualitatively by visual inspection of segmentation outputs across complete assay images. Fig. 2C,D illustrate representative examples of segmentation masks at different stages of the pipeline, including the initial candidate masks produced by the segmentation model and the refined masks obtained after iterative filtering.

Across assay images containing approximately 200 leaves arranged on a uniform background, the pipeline successfully isolated individual leaves with minimal fragmentation or merging. The iterative refinement procedure effectively adds new leaf-masks in the segmentation results, as seen by the yellow arrow in Fig. 2. Visual inspection confirmed that the final segmentation masks corresponded closely to individual physical leaves and were suitable for downstream lesion analysis.

### Lesion estimation accuracy

The accuracy of lesion area estimation was evaluated on the held-out test set using MAPE and MAE. Fig. 6 shows the relationship between predicted lesion areas and manually annotated ground-truth values. On the test set, the model achieved a MAPE of 12.9%. This result indicates that the lesion estimation model produces lesion area measurements that are consistent with manually annotated reference values across a range of lesion sizes.

**Fig 6.**
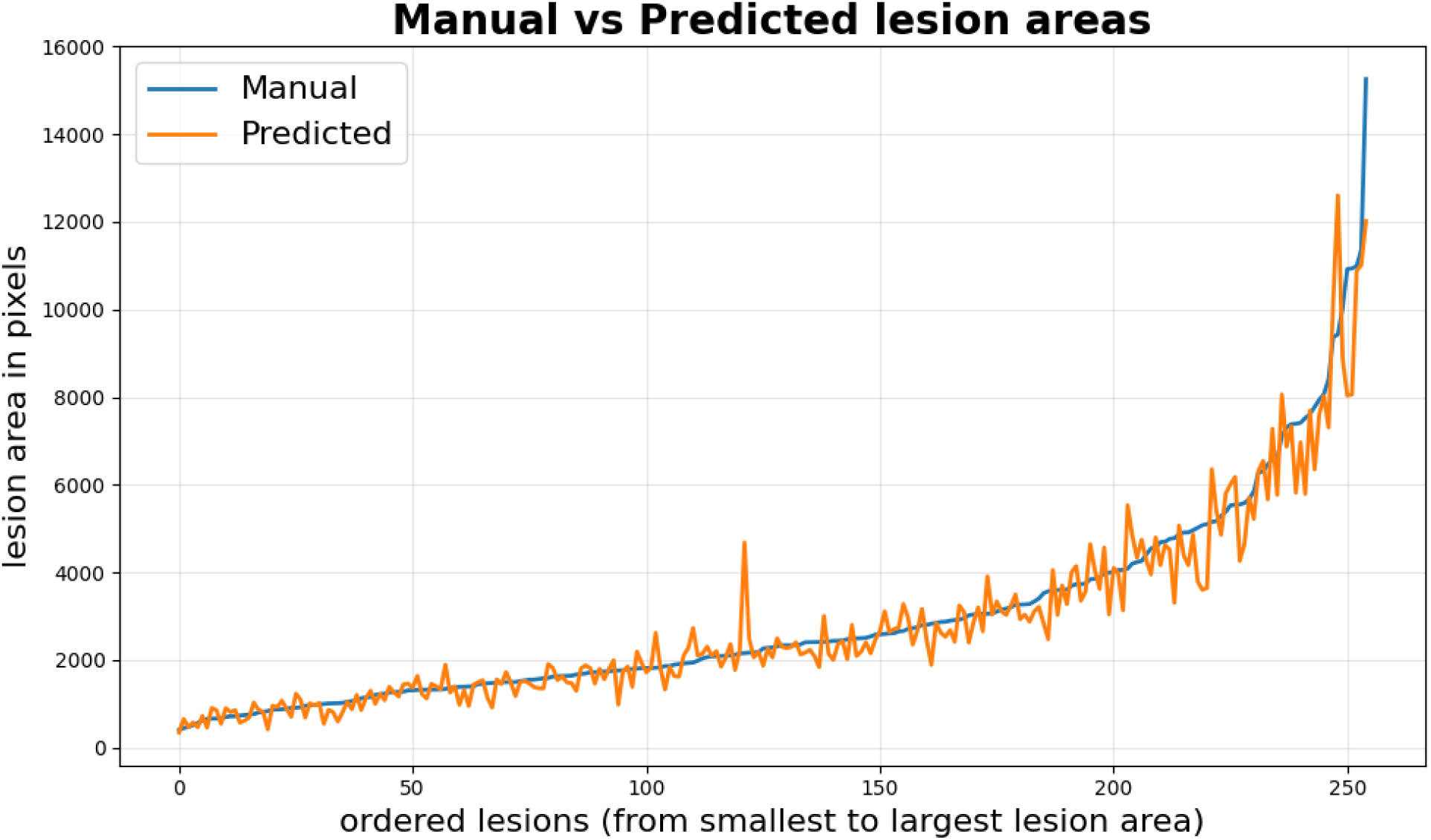
Predicted versus manual lesion areas. NASNet A test-set performance shown as predicted lesion area versus manually annotated lesion area.

### Performance across lesion-size groups

Model performance was evaluated separately for small, intermediate, and large lesion-size groups to characterize behavior across disease severity levels. Prediction accuracy was assessed using MAE, which reflects absolute deviation in pixel units, and MAPE, which normalizes error relative to lesion size. Quantitative results are summarized in Table 2, and performance trends are visualized in Fig. 7.

**Table 2.** Prediction accuracy by lesion-size group. Results for the deep learning–based model and color-based baselines are reported as mean absolute error (MAE, pixels) and mean absolute percentage error (MAPE, %).

| Group | Model | MAE | MAPE (%) |
| --- | --- | --- | --- |
| Small | NASNet A | 155.25 | 14.90 |
|  | HSV Threshold | 1644.37 | 167.32 |
|  | K-means | 5644.25 | 553.09 |
| Intermediate | NASNet A | 275.08 | 12.34 |
|  | HSV Threshold | 2579.25 | 116.80 |
|  | K-means | 6171.49 | 284.69 |
| Large | NASNet A | 652.14 | 11.56 |
|  | HSV Threshold | 3210.94 | 64.75 |
|  | K-means | 6132.74 | 130.15 |
| Overall | NASNet A | 360.83 | 12.93 |
|  | HSV Threshold | 2478.19 | 116.29 |
|  | K-means | 5982.82 | 322.64 |

**Fig 7.**
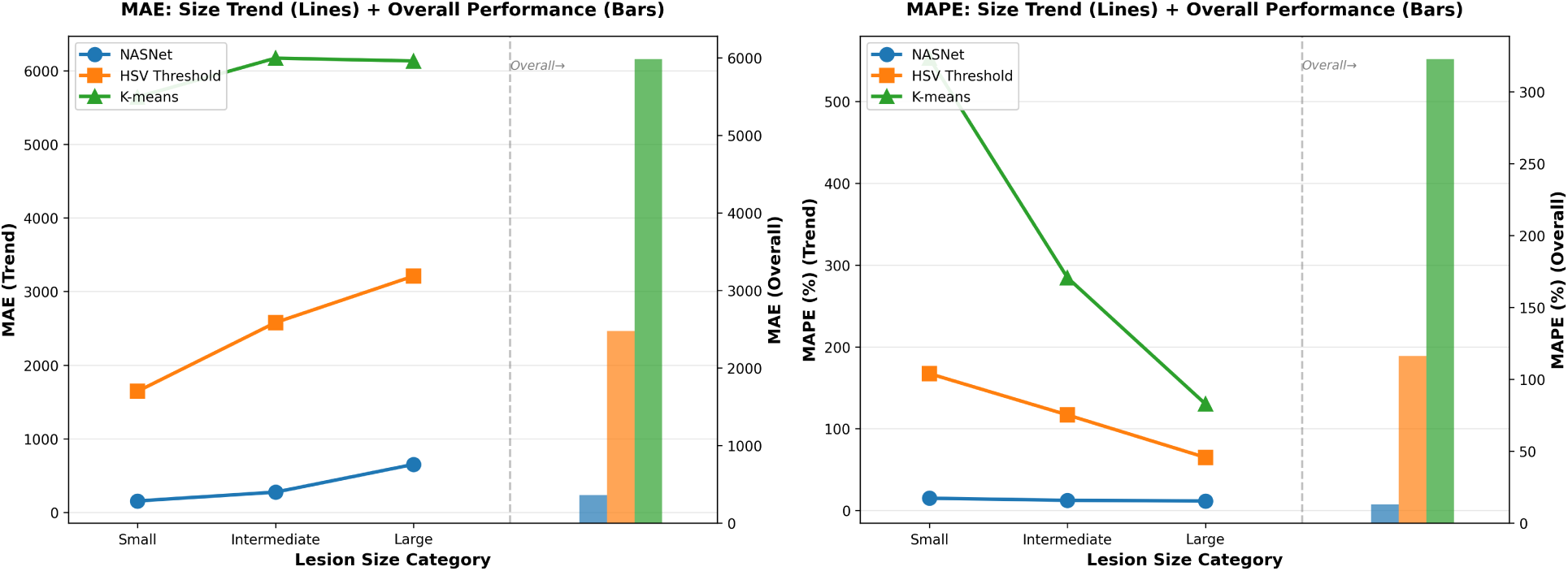
Prediction accuracy across lesion-size categories. Line plots show MAE (left) and MAPE (right) trends for small, intermediate, and large lesion groups, while bar plots indicate overall performance on the full test set. The NASNet A-based model maintains consistently lower error across lesion sizes than the HSV thresholding and K-means clustering baselines.

Across all lesion-size groups, the NASNet A-based model consistently achieved substantially lower MAE and MAPE values than both color-based baselines. For the NASNet-based model, MAE increased with lesion size, reflecting the larger absolute lesion areas in the intermediate and large groups. In contrast, MAPE decreased slightly as lesion size increased, indicating that relative prediction error was more pronounced for small lesions than for larger ones.

The color-based baselines exhibited markedly different trends. Both HSV thresholding and K-means clustering produced very large MAE values across all lesion-size groups, indicating substantial absolute overestimation of lesion area. These errors were especially severe for small lesions, where MAPE exceeded 100%, reflecting the sensitivity of relative error metrics to small denominators. While MAPE values for the baselines decreased for larger lesions, absolute errors remained high, suggesting that these methods consistently misclassified healthy tissue as lesion regardless of disease severity.

Overall performance on the full test set followed similar patterns. The NASNet A-based model maintained stable relative accuracy across lesion sizes, achieving an overall MAPE of 12.9%, whereas both color-based baselines exhibited high absolute and relative errors and strong dependence on lesion size.

Representative qualitative examples illustrating model behavior across lesion-size groups are shown in Fig. 9. These examples highlight the ability of the NASNet A-based model to adapt to varying lesion extents, while color-based methods tend to overestimate lesion regions, particularly for leaves with small lesion sizes.

### Consistency and throughput comparison

To assess the consistency of manual lesion scoring, a subset of 12 leaf images was randomly selected from the dataset and independently rescored by the same biologist. Fig. 8 demonstrates those rescored leaves and the evaluated lesion areas in pixels. Agreement between the original scores and the rescored values was quantified using MAPE.

**Fig 8.**
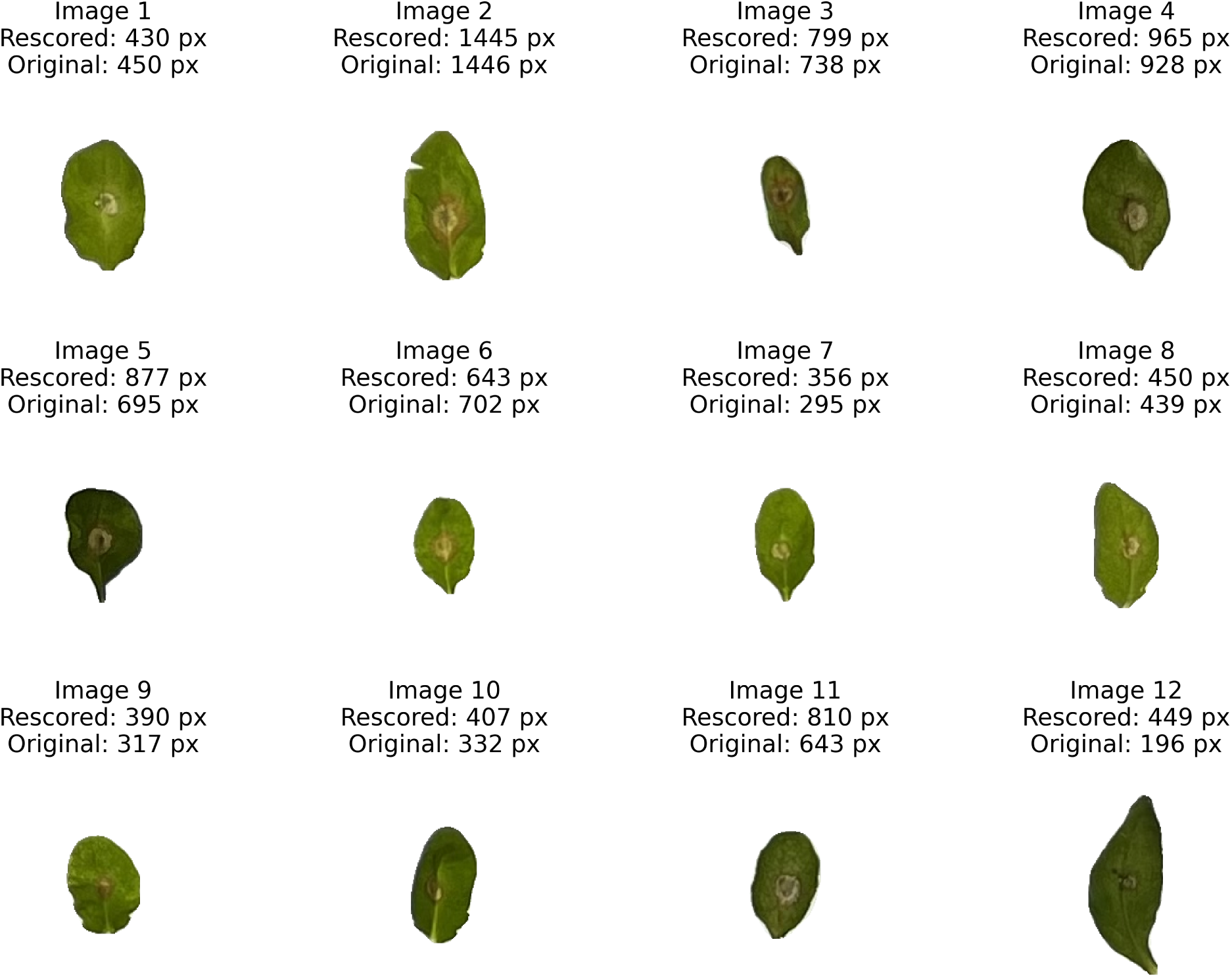
Repeated manual scoring of representative leaves. Consistency of manual lesion scoring based on repeated annotation of 12 randomly selected leaf images by the same biologist. Lesion area is reported in pixel units.

Across the 12 samples, the rescoring exhibited a mean absolute error of 83.3 pixels and a mean absolute percentage error of 10%. This intrarater error is close in magnitude to the overall LIME test-set MAPE of 12.9% reported in Table 2, which helps contextualize model performance relative to human repeatability. These results indicate non-negligible intrarater variability in manual lesion quantification, particularly for leaves with smaller lesion areas.

Runtime performance was assessed by measuring the total time required to process a complete assay image containing approximately 200 leaves. Under GPU-accelerated execution, the full LIME pipeline—including segmentation, lesion area estimation, and post-processing—required approximately 15 minutes per assay. In contrast, manual lesion scoring required approximately 60 seconds per leaf, corresponding to a total processing time of roughly 3 hours for an equivalent assay. Based on these measurements, LIME provides an approximate 13-fold reduction in processing time compared to manual scoring for large-scale assays. Importantly, while the intrarater comparison was based on 12 leaves and does not directly quantify long-session fatigue effects, automated inference avoids fatigue- and attention-related drift that can affect repetitive manual visual scoring tasks as workload increases.

## Discussion

### Interpretation of accuracy, lesion-size effects, and human variability

Quantitative evaluation of lesion area estimation demonstrated that the NASNet A-based model achieved a MAPE of approximately 12.9% on the held-out test set. Stratified analysis across lesion-size groups further showed that prediction accuracy remained relatively stable for small, intermediate, and large lesions, with MAPE values ranging from 11.6% to 14.9%. In contrast, both color-based baselines exhibited substantially higher error and pronounced sensitivity to lesion size, particularly for small and intermediate lesions.

These results indicate that the deep learning–based model generalizes more consistently across disease severity levels than threshold- or clustering-based approaches. The group-wise comparison highlights that color-based methods tend to overestimate lesion area when lesion appearance overlaps with healthy tissue coloration, an effect that becomes more pronounced for smaller lesions where absolute area differences translate into large relative errors.

The intrarater rescoring indicates non-negligible variability even when annotations are performed by the same expert. This variability underscores the inherent subjectivity of manual lesion quantification and supports interpreting model error as agreement with human reference measurements rather than deviation from an objective ground truth. Within this context, the accuracy achieved by the NASNet A-based model falls within the range of variability observed in human scoring.

### Practical implications for phenotyping pipelines

Beyond predictive accuracy, the results demonstrate the practical advantages of a fully automated lesion quantification pipeline for large-scale leaf phenotyping studies. By eliminating manual threshold selection and subjective visual scoring, LIME reduces operator-dependent variability and ensures consistent measurement across experiments. This is particularly important in comparative phenotyping settings, where subtle differences in disease severity must be evaluated across large numbers of samples.

The manual intrarater analysis (10% MAPE on 12 leaves) and the LIME test-set error (12.9% MAPE) indicate comparable accuracy in a small-scale repeated-scoring setting. However, the practical advantage of machine learning becomes stronger with larger cohorts: model performance is stable across throughput, whereas manual scoring quality can degrade during prolonged repetitive annotation due to human workload and attentional effects. In this sense, automation provides not only speed, but also consistency preservation as sample counts increase.

Throughput analysis further highlights the scalability of the approach. Processing a complete assay image containing approximately 200 leaves required approximately 15 minutes under GPU-accelerated execution, compared to roughly 3 hours for manual scoring. This reduction in analysis time enables rapid iteration over large experimental cohorts and makes quantitative lesion analysis feasible in high-throughput experimental designs, such as genetic screens or multi-condition studies.

The deterministic nature of the pipeline also ensures reproducibility and consistency compared with manual scoring: repeated analysis of the same assay image yields identical results, facilitating comparisons across research groups.

### Limitations and future work

Despite its overall robustness, several limitations were observed. Segmentation performance can degrade in cases where leaves overlap extensively or when background artifacts exhibit color characteristics similar to leaf tissue. Such errors may propagate to downstream lesion estimation, emphasizing the importance of standardized assay layout and imaging conditions.

Additionally, lesion area estimation is formulated as a regression problem that predicts aggregate lesion area without explicitly delineating lesion boundaries. While this approach avoids the ambiguity associated with pixel-level lesion annotation, it does not provide spatial localization of lesions within individual leaves. Meanwhile, the occlusion map shown in Fig. 9 only indicates roughly which regions most influence the NASNet A prediction, rather than explicit lesion boundaries. Consequently, the current implementation is best suited for applications where total lesion burden is the primary metric of interest rather than detailed lesion morphology.

**Fig 9.**
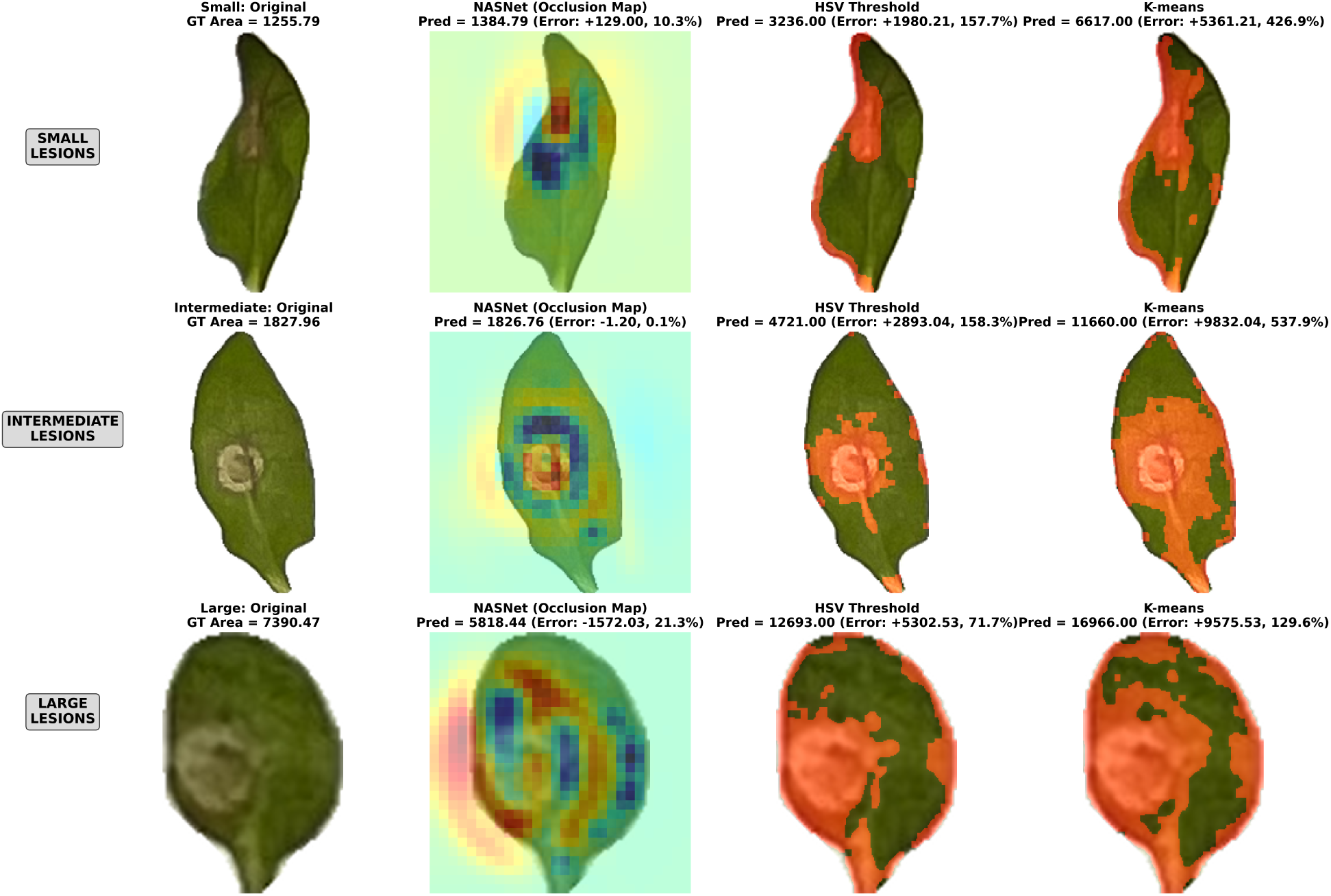
Qualitative examples across lesion-size categories. Each row corresponds to a lesion-size group (small, intermediate, large) and shows, from left to right, the original leaf image, the NASNet A occlusion map, the lesion regions identified by HSV thresholding, and the lesion regions identified by K-means clustering. Occlusion maps indicate regions that most influence the NASNet A prediction rather than explicit lesion boundaries.

Future work will focus on extending the pipeline to additional pathosystems and symptom types through retraining of the lesion estimation model on new datasets. Further improvements may also include enhanced handling of complex assay layouts and the incorporation of alternative lesion quantification strategies when spatial lesion information is required.

## Conclusion

In this study, we presented LIME, a fully automated pipeline for high-throughput quantification of leaf lesion severity from assay images. By integrating zero-shot leaf segmentation with a deep learning–based lesion area estimation model, LIME enables objective and reproducible measurement of plant disease severity across large-scale phenotyping experiments.

Comprehensive evaluation on *Arabidopsis thaliana* leaves infected with *Sclerotinia sclerotiorum* demonstrated that the proposed approach achieves consistent prediction accuracy across lesion-size groups and outperforms color-based baselines. Model accuracy was comparable to observed intrarater variability in manual scoring, while reducing analysis time by an order of magnitude relative to human annotation. These results indicate that LIME provides a practical, scalable, and reliable solution for quantitative lesion analysis in standardized plant pathology assays.

## Acknowledgments

Special thanks to the Terrabyte group at the University of Winnipeg and the University of Manitoba for their ideas and guidance in creating this work.

